# Robust Estimation of 1/f Activity Improves Oscillatory Burst Detection

**DOI:** 10.1101/2022.03.24.485674

**Authors:** Robert A. Seymour, Nicholas Alexander, Eleanor A. Maguire

## Abstract

Neural oscillations often occur as transient bursts with variable amplitude and frequency dynamics. Quantifying these effects is important for understanding brain-behaviour relationships, especially in continuous datasets. To robustly measure bursts, rhythmical periods of oscillatory activity must be separated from arrhythmical background 1/f activity, which is ubiquitous in electrophysiological recordings. The Better OSCillation (BOSC) framework achieves this by defining a power threshold above the estimated background 1/f activity, combined with a duration threshold. Here we introduce a modification to this approach called fBOSC which uses a spectral parametrisation tool to accurately model background 1/f activity in neural data. fBOSC (which is openly available as a MATLAB toolbox) is robust to power spectra with oscillatory peaks and can also model non-linear spectra. Through a series of simulations, we show that fBOSC more accurately models the 1/f power spectrum compared with existing methods. fBOSC was especially beneficial where power spectra contained a “knee” below ∼0.5-10 Hz, which is typical in neural data. We also found that, unlike other methods, fBOSC was unaffected by oscillatory peaks in the neural power spectrum. Moreover, by robustly modelling background 1/f activity, the sensitivity for detecting oscillatory bursts was standardised across frequencies (e.g. theta- and alpha-bands). Finally, using openly available resting state magnetoencephalography and intracranial electrophysiology datasets, we demonstrate the application of fBOSC for oscillatory burst detection in the theta-band. These simulations and empirical analyses highlight the value of fBOSC in detecting oscillatory bursts, including in datasets that are long and continuous with no distinct experimental trials.

**GRAPHICAL ABSRACT:** 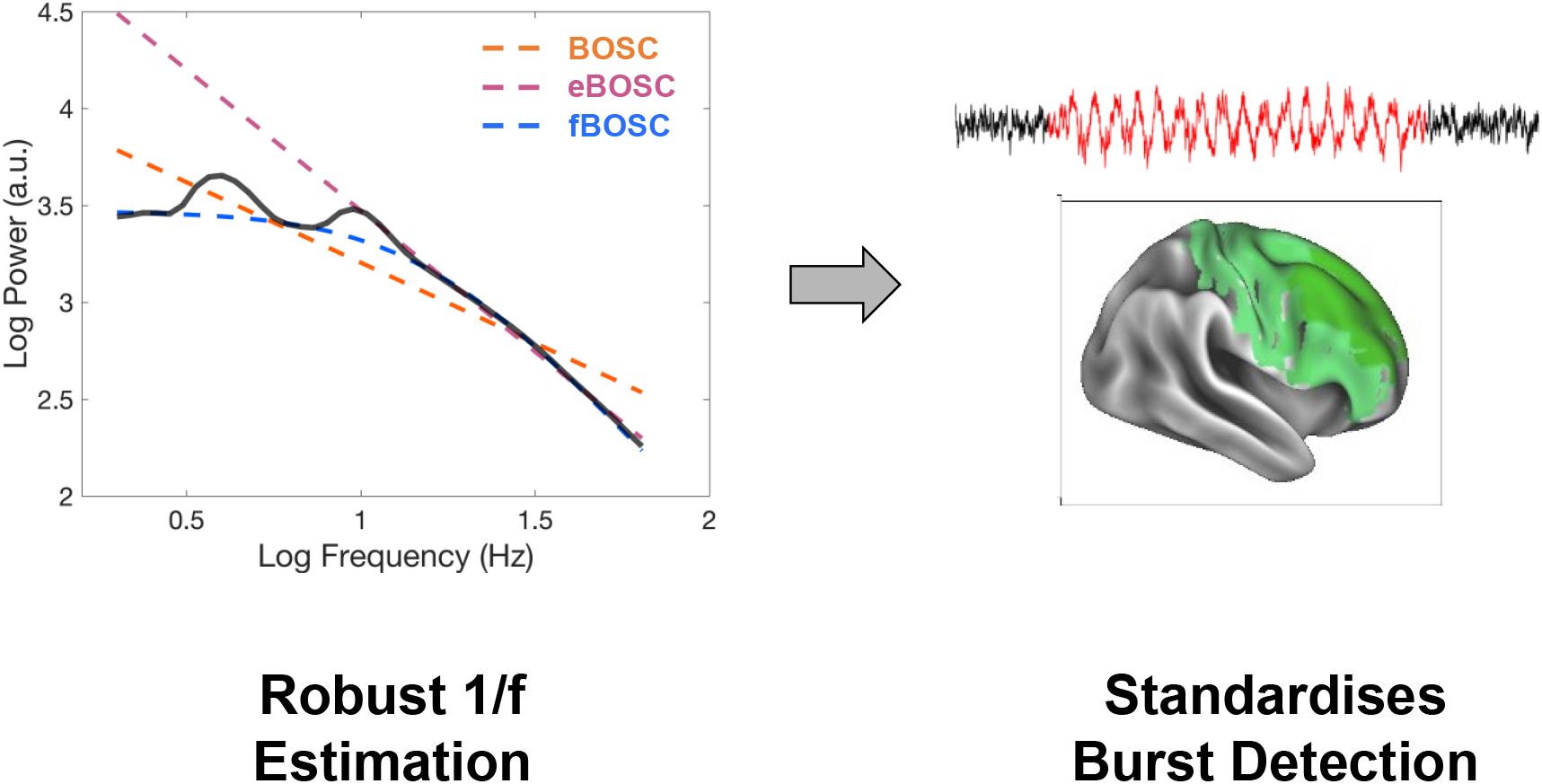

To determine a power threshold for burst detection, the Better OSCillation framework (BOSC) estimates background 1/f activity by modelling neural power spectra. Here we introduce a modification, termed fBOSC, to more robustly estimate 1/f activity in situations with prominent oscillatory peaks and/or the presence of a non-linear “knee” in the power spectrum. This was shown to standardise burst detection across frequency bands in both simulated and empirical data.

## INTRODUCTION

Rhythmical signals are present in many different types of neural data, from single neuron measurements to non-invasive magnetoencephalography (MEG) or electroencephalography (EEG) recordings of large-scale population dynamics (Başar et al., 2000; Buzsaki, 2006). Changes in the amplitude of these neural oscillations are linked to a range of cognitive tasks within specific frequency bands. For example, motor tasks are associated with dynamic beta-band (13-30 Hz) changes (Barratt et al., 2018; Neuper & Pfurtscheller, 2001), and higher-level executive functions (e.g. working memory, memory encoding) are associated with theta-band (3-7 Hz) changes (Costers et al., 2020; Herweg et al., 2020; Roux & Uhlhaas, 2014). From a theoretical perspective, oscillations have been argued to play a mechanistic role in the dynamic temporal and spatial organisation of neural activity (Bastos et al., 2015; Buzsaki, 2006; Fries, 2015). Furthermore, perturbations to oscillations are associated with several clinical conditions, including Autism spectrum disorder (Kessler et al., 2016; Seymour et al., 2019), schizophrenia (Kirihara et al., 2012; Thuné et al., 2016) and mild traumatic brain injury (Allen et al., 2021).

Electrophysiological analyses typically average oscillatory power across trials. However, oscillations within single trials are often high-amplitude and transient, occurring as short “bursts” of activity (Jones, 2016; Jones et al., 2009; Stokes & Spaak, 2016). The characteristics of this bursting behaviour often go unstudied, meaning that potentially important information in neural datasets is missed. Single trial burst analyses allow the separation of rhythmical burst amplitude from duration (Kosciessa et al., 2020; Quinn et al., 2019). Furthermore, characterising oscillations as bursts rather than sustained rhythmical signals is more physiologically faithful, which has implications for brain-behaviour relationships (Stokes & Spaak, 2016). For example, within the motor literature, it has recently been demonstrated that the rate and timing of beta bursts in humans is more predictive of behaviour than mean beta amplitude (Bonaiuto et al., 2021; Jana et al., 2020). Consequently, there is increasing interest in characterising the bursting properties of neural oscillations (Bonaiuto et al., 2021; Jones, 2016; Kosciessa et al., 2020; Stokes & Spaak, 2016). This will be particularly important for continuous datasets which cannot be split into distinct trials, such as naturalistic paradigms involving free movement (Seymour et al., 2021; Stangl et al., 2021).

Rhythmical bursts occur in the context of a continuous, arrhythmic, fractal-like background component in neural data (He, 2014; He et al., 2010; Miller et al., 2009). The power spectral density of this component decreases logarithmically with frequency, such that P ≈1/f, where β is the power-law exponent (Chaudhuri et al., 2017; Miller et al., 2009). β typically varies between 1 and 3 in human electrophysiological data (He, 2014; He, 2010; Miller et al., 2009). Here we refer to this activity as “(background) 1/f activity”, but others call it “scale-free” (He et al., 2010) or “aperiodic” (Donoghue, et al., 2020b). This type of activity is ubiquitous across a wide range of systems, from empty-room EEG and MEG recordings to economic data (He et al., 2010). In the context of brain dynamics, neural 1/f activity is not simply a summation of multiple oscillations, but potentially reflects the population firing rate (Manning et al., 2009; Miller, 2010), and also relates to the balance between excitatory and inhibitory neural circuits (Gao et al., 2017). Dynamic changes to background 1/f activity have also been associated with various aspects of cognitive function independently of oscillations (Gao et al., 2020; Helfrich et al., 2021; Lendner et al., 2020; Wilson et al., 2022). Given their different neural origins, it is vital to robustly separate background 1/f activity from oscillations (Donoghue et al., 2020a; Donoghue et al., 2020b; Gerster et al., 2021; He, 2014). This presents a challenge from a data analysis perspective, given that electrophysiological tools concurrently measure both rhythmic and arrhythmical signals (Donoghue et al., 2021).

There are several burst detection methods that aim to solve this problem, but here we focussed on the Better OSCillation (BOSC) framework (Hughes et al., 2012; Whitten et al., 2011) which works as follows: for each frequency of interest, a power threshold is defined based on the modelled background 1/f spectrum. In addition, a duration threshold is defined, typically equivalent to 2-3 oscillatory cycles. An oscillatory burst is said to be detected when both the power and duration thresholds are exceeded. BOSC has been applied successfully to a range of electrophysiological datasets in humans and rodents (Caplan et al., 2001; Hughes et al., 2012; Stangl et al., 2021; Whitten et al., 2011).

Here, we introduce a modification to the BOSC method by modelling the background 1/f activity with a recently developed spectral parametrisation tool (Donoghue et al., 2020b). Using simulated data, we show that our tool, called fBOSC, recovers 1/f signals with greater accuracy than existing methods, especially in situations with non-linear power spectra. Simulations also show that by modelling the 1/f spectrum more accurately, fBOSC standardises burst detection across frequency bands. Finally, we demonstrate the empirical utility of fBOSC using openly available MEG and intracranial EEG (iEEG) datasets.

## MATERIALS AND METHODS

### The BOSC framework and different approaches to 1/f fitting

The BOSC framework determines the frequency and duration of transient oscillatory bursts above and beyond the background 1/f power spectrum present in neurophysiological data (Hughes et al., 2012; Whitten et al., 2011). The first step involves a time-frequency decomposition of each experimental trial, typically using Morlet wavelets to control time-frequency trade-offs. The logarithmically transformed power spectrum, averaged over trials, is then used to fit the background 1/f activity over all frequencies of interest (e.g. 2-64 Hz). A power threshold based on this 1/f fit is then calculated, for example, at the 99th percentile of the theoretical probability distribution (with chi-squared form) of power values at a given frequency. A duration threshold is also defined, typically equivalent to 2 or 3 oscillatory cycles. For each trial, if the power at a given frequency exceeds the threshold and lasts longer than the duration threshold, an oscillation is said to be detected.

A key part of the BOSC framework is the accurate modelling of 1/f background activity, as it is used to directly define the power threshold. The original BOSC implementation (Hughes et al., 2012; Whitten et al., 2011) uses ordinary least squares regression to fit the power spectrum in log-log space (Figure 1 left). However, when a prominent peak(s) exists in the power spectrum, this approach can lead to a skewed 1/f fit. For example, Whitten et al., (2011) noted that performing the background fit on data with a large alpha-band peak (where eyes were closed) led to different results compared to data with a relatively flat spectrum (when eyes were open). To address this, an extended BOSC (eBOSC) implementation (Kosciessa et al., 2020) was introduced using MATLAB’s robustfit function to down-weight outliers. In addition, eBOSC allows the user to exclude certain peak frequencies during the 1/f fit (see Figure 1, middle).

**FIGURE 1.**
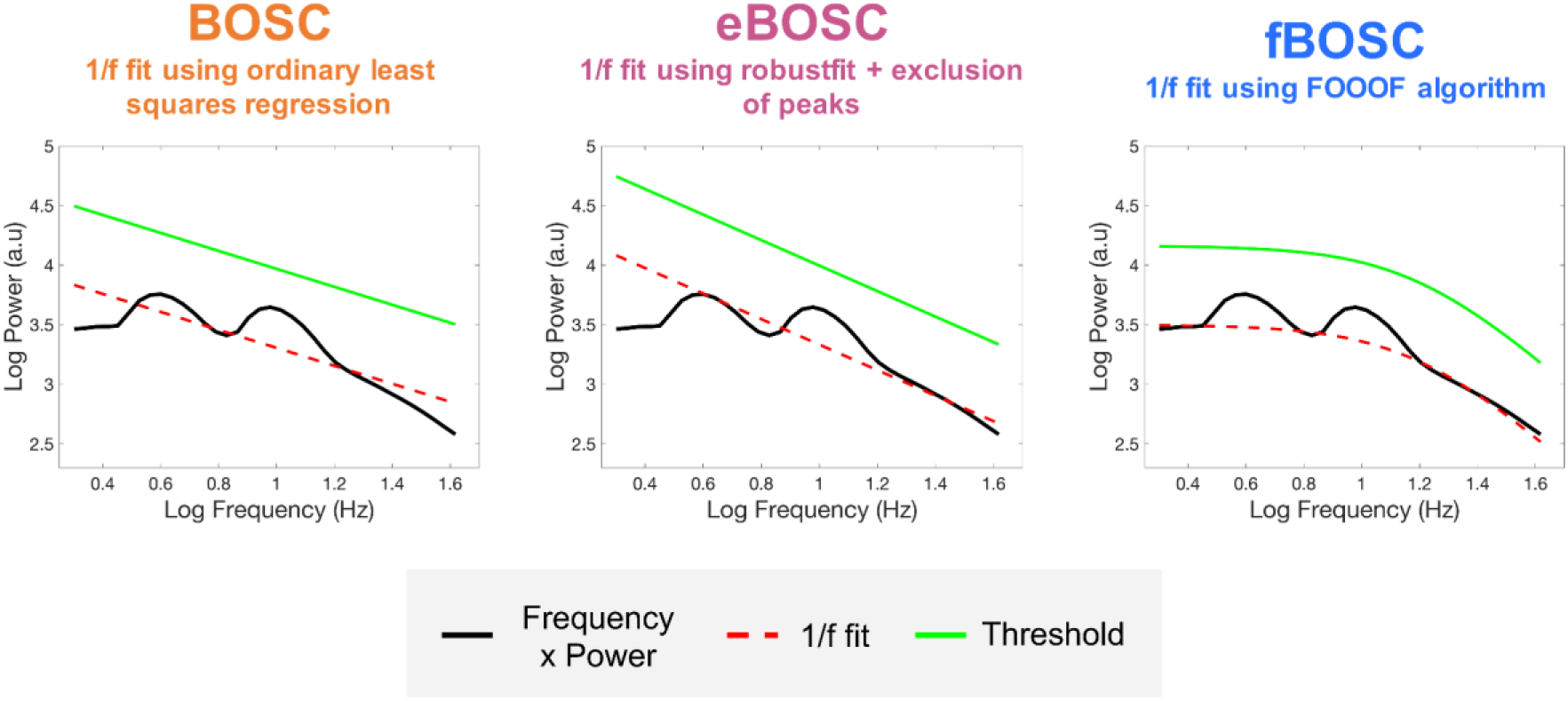
Different methods for modelling the background 1/f power spectrum within the BOSC framework. The original BOSC implementation uses an ordinary least squares approach. The eBOSC toolbox uses MATLAB’s robustfit function, whilst also allowing the user to exclude peaks. fBOSC uses the FOOOF algorithm to parametrise the neural spectra. In this example, a non-linear 1/f frequency spectrum with a prominent knee is plotted in log-log space. Unlike fBOSC, the 1/f fits from BOSC and eBOSC (red dotted line) fail to model this non-linear spectrum. Power thresholds (plotted in green) are therefore higher for frequencies below the knee when using BOSC and eBOSC compared to fBOSC.

However, two issues remain. First, where oscillatory peaks exist in the power spectrum, the eBOSC approach requires manual specification of peak frequencies, which is not ideal for data automation and processing pipelines. Furthermore, exclusion of multiple frequencies in the power spectrum may lead to a poor 1/f fit. Second, where power spectra are non-linear when plotted in log-log space, the linear methods used by BOSC and eBOSC will be unsuitable. This is especially problematic during the analysis of lower frequency oscillatory bursts (e.g. delta- and theta-bands), as human neurophysiological data often exhibits a bend or “knee” at ∼0.5-10 Hz (He, 2014; He et al., 2010). Furthermore, there is emerging evidence that the broadband power spectrum of humans is highly non-linear, and is best modelled by the sum of two Lorentzian functions (Chaudhuri et al., 2017; Gao et al., 2017).

To address these issues, we introduce a modification to the 1/f fitting procedure under the BOSC framework, using the “fitting oscillations and one over f” (FOOOF) spectral parametrisation algorithm (Donoghue et al., 2020b). We call this modification fBOSC (FOOOF+BOSC, see Figure 1 right). In short, the FOOOF algorithm performs an initial 1/f fit and iteratively models oscillatory peaks above this background as gaussians. These peaks are removed from the spectra and the 1/f fit is performed again. Consequently, the fit is not influenced by oscillatory peaks (Donoghue et al., 2020b; Wilson et al., 2022), and no manual selection of peak frequencies is required, as is the case for eBOSC. FOOOF can also model the 1/f power spectrum as either linear or non-linear. In the case of latter, a variable knee parameter is included. This addresses the second issue with the BOSC/eBOSC fitting approaches.

An additional advantage of FOOOF is that the 1/f fit returns several parameters describing the shape of the spectrum over frequencies of interest (*f*). First, the offset parameter (*b*) which corresponds to the overall up/down translation of the whole spectrum. Second, the exponent (*β*) describes the rate of decay over frequency (Chaudhuri et al., 2017; Miller et al., 2009). The third optional parameter, *k*, describes the knee in the power spectrum. These parameters are estimated by the fit of the function (*L*) to the background 1/f activity, expressed as:

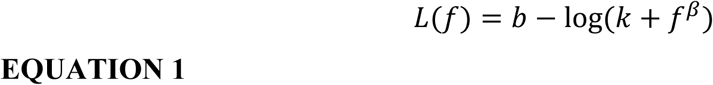

### Code availability

We share the MATLAB code for fBOSC openly at https://github.com/neurofractal/fBOSC. The code is built on top of the original BOSC (Caplan et al., 2001) and eBOSC (Kosciessa et al., 2020) code and is shared under a GNU General Public License v3.0. fBOSC is also compatible with the post-processing options included with eBOSC. For all results presented here we used v0.1 of fBOSC and the original Python implementation of FOOOF (Donoghue et al., 2020b). However a MATLAB version of the same algorithm has also been implemented within fBOSC, adapted from Brainstorm (Tadel et al., 2011). All FOOOF parameters were set to the Python defaults: smallest peak width = 0.5; largest peak width =12.0; maximum number of peaks = infinite; minimum peak height = 0; relative threshold for detecting peaks = 2 standard deviations.

### Simulations

Various simulations were performed to assess the performance of BOSC, eBOSC and fBOSC. We generated two types of data to mimic background, aperiodic neural activity with either linear or non-linear 1/f slopes when plotted in log-log space. Data with linear 1/f power spectra were generated by filtering randomly generated data using the cnoise MATLAB toolbox (https://people.sc.fsu.edu/~jburkardt/m_src/cnoise/cnoise.html). The β exponent was equal to 2, to approximately mimic human neurophysiological data. Data with non-linear 1/f power spectra were generated by convolving Poisson activity with exponential kernels that mimic the shape of post-synaptic potentials, using the Python neurodsp toolbox (`sim_synaptic_current` function; Cole et al., 2019). These neurophysiologically plausible data had a prominent knee in the power spectrum when plotted in log-log space. For both linear and non-linear simulations, two hundred trials, each lasting 60 s, were computed at a 500 Hz sampling rate. Sine waves at 4 Hz or 10 Hz were then added to simulate oscillatory bursts in the theta-band alone or the alpha-band alone. These lasted 15 s in total, but were broken into transient bursts and placed pseudo-randomly (separated by at least 0.5s) within the 60 s-long trial. To approximately mimic physiological bursting behaviour, each sine wave varied randomly between 2-7 cycles (theta-band) or 8-30 cycles (alpha-band), and were separated by a minimum of 0.5 s (Aghajan et al., 2017; Kosciessa et al., 2020; Whitten et al., 2011). The amplitude of the sine waves were equivalently scaled to specific signal-to-noise ratios (SNRs) based on the relative amplitude of band-pass filtered data (1 Hz bandwidth) at 4 Hz or 10 Hz. Note that SNR in the context of our study refers to the *relative* measure between background 1/f power and rhythmical power at 4 Hz or 10 Hz, rather than the overall SNR across all frequencies (Kosciessa et al., 2020). To approximate the variability of theta/alpha bursts in neurophysiological data, the SNR of each burst was varied randomly between 5-12. Finally, we simulated data with oscillatory bursts at both theta and alpha frequencies by concatenating 60 s-long trials together (i.e. creating 120 s-long trials, with theta bursts in the first 60 s and alpha bursts in the last 60 s).

For BOSC, eBOSC and fBOSC analyses, we used frequency sampling at logarithmically-spaced frequencies (log base 2) between 2 to 64 Hz. Time-frequency analysis was conducted using Morlet wavelets (6-cycles). This was performed on the simulated data before and after the oscillatory bursts were added. The mean power spectrum over all trials was then used for the 1/f fit in BOSC, eBOSC and fBOSC. For eBOSC, frequencies between 3-7 Hz (theta) and 7-13 Hz (alpha) were excluded from the 1/f fit. For fBOSC, the aperiodic mode was modelled with a knee for the non-linear 1/f data, and without a knee (fixed) for the linear 1/f data. For each trial, the 1/f fit from all three methods was compared with the power spectrum of the original 1/f data using the root mean squared error (RMSE) metric.

When FOOOF is used to model power spectra with a knee, the 1/f fit will contain an extra term, *k* (see Equation 1), corresponding to a bend in the power spectrum when plotted in log-log space. Consequently, for the non-linear simulations, the 1/f fit from fBOSC will be more complex than for eBOSC and BOSC. To rule out the possibility that RMSE differences between methods were influenced by differences in model complexity (i.e. over-fitting) we conducted two follow-up analyses. First, we performed 10-fold cross validation. The simulated data were split into a training dataset (90% of trials) and a testing dataset (10% of trials). Using the training data, the background 1/f activity was estimated using BOSC, eBOSC and fBOSC and the resulting 1/f model fit from each method was used to predict the 1/f power spectrum in the testing dataset. This was repeated iteratively for each fold of the data. Second, we repeated the simulation analyses using the non-linear 1/f data and the Akaike Information Criterion (AIC) rather than RMSE to quantify the error between the ground truth and predicted 1/f fit. AIC is defined in Equation 2, where *n* is the number of observations, *SSE* is the sum of square error and *c* is the number of independently adjusted parameters within the model. This is a commonly used method for model selection since the *c* term penalises models for complexity.

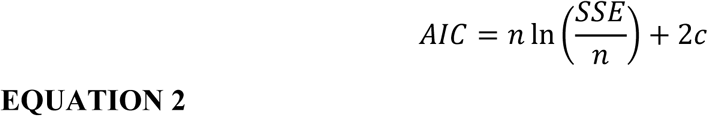

In a separate analysis, using BOSC, eBOSC and fBOSC, we used the 1/f fitting procedures to define a power threshold (0.99, chi-squared distribution), combined with a duration threshold (3 oscillatory cycles) to detect oscillatory bursts. For each trial, the hit rate (time points where a burst was simulated and detected) and the false alarm rate (time points where a burst was not simulated but detected) was quantified for the theta- and alpha-bands.

### Resting state MEG data analysis

We compared burst detection between BOSC, eBOSC and fBOSC using an openly available MEG dataset from the Young Adult Human Connectome Project (HCP; Larson-Prior et al., 2013). Resting state data from the first 50 participants were used (26 female, 24 male; participant numbers 100307 to 214524; first run only) which had been pre-processed using a standardised HCP MEG Pipeline (see: https://www.humanconnectome.org/software/hcp-meg-pipelines). The resting state scan involved participants looking at a screen and fixating on a red cross for 6 minutes. Sensor-level data were mapped to source space using a linearly constrained minimum variance beamformer (Van Veen et al., 1997), as implemented in the Fieldtrip toolbox (Oostenveld et al., 2011). For the forward model the participant’s T1-weighted structural MRI scan was used to create a single-shell description of the inner surface of the skull (Nolte, 2003). Using SPM12, a non-linear spatial normalisation procedure was used to construct a volumetric grid (8 mm resolution) registered to the canonical Montreal Neurological Institute brain. The source-localised data were then parcellated into 42 cortical regions of interest (ROIs) based on a down-sampled version of the whole brain HCP multimodal atlas (Glasser et al., 2016). The right dorsolateral prefrontal cortex had the highest abundance of theta bursts, and we therefore concentrated further analyses on this ROI. For burst detection, the power threshold for rhythmicity at each frequency was set at the 99^th^ percentile of a chi-squared distribution of power values, in combination with a duration threshold of 3 oscillatory cycles.

### Resting state iEEG data analysis

We also analysed an openly available resting state iEEG dataset, originally published by Miller et al. (2012, 2017). The data were collected from 10 patients (aged 18-42) with implanted electrocorticographic grids for the monitoring and treatment of medically-refractory epilepsy. A total of 533 electrodes were analysed across the 10 patients, which were mainly located across bilateral frontal and frontotemporal areas (exact electrode placements were not provided). Patients were instructed to fixate on an ‘X’ which was located on a wall 3 m away, for a duration of 2-3 minutes. Data were sampled at a rate of 1000 Hz. Raw iEEG timeseries were loaded into MATLAB, down-sampled to 200 Hz, notch filtered at 60 Hz, followed by a high-pass filter at 0.5 Hz and low-pass filter at 90 Hz (5th order Butterworth filters applied bidirectionally to achieve zero-phase shift). All patients participated in a purely voluntary manner, after providing informed written consent, under experimental protocols approved by the Institutional Review Board (IRB) of the University of Washington (#12193). All patient data were anonymised according to IRB protocol, in accordance with HIPAA mandate.

### Statistical analysis

Statistical analyses were conducted using the JASP software package (Love et al., 2019). One-way ANOVAs were used to compare RMSE, hit rate, false alarm rate and theta abundance across the three methods BOSC, eBOSC and fBOSC. Post-hoc tests were then conducted with Bonferroni correction. For the knee versus fixed analysis, a paired two-tailed t-test was used.

## RESULTS

### fBOSC reduces 1/f fit error

To compare our fBOSC method, which combines FOOOF spectral parametrisation with the BOSC framework, to existing methods (BOSC and eBOSC, see Figure 1) we performed a series of simulations. Data with a linear or non-linear background 1/f power spectrum were combined with simulated oscillatory bursts in the theta-band (4 Hz), the alpha-band (10 Hz), or both bands, at SNRs randomly varying between 5-12. The RMSE between the simulated 1/f power spectrum (with known ground truth) and the 1/f fit using BOSC, eBOSC and fBOSC was calculated. Note that lower RMSE values indicate better performance. We chose to simulate sinusoids at 4 Hz and 10 Hz as we expected these frequency bands to show the greatest differences between the methods, especially when embedded within a non-linear 1/f power spectrum.

Results for the simulated data with a linear 1/f power spectrum are shown in Figure 2A-C. Across all three simulated oscillatory burst conditions (theta alone, alpha alone, theta and alpha) there was a main effect of method, F(2,597) > 1057, p < 0.001. Follow-up tests showed that fBOSC outperformed BOSC, with significantly lower RMSE values across all bursting conditions, all p < 0.001. eBOSC had lower RMSE values than BOSC, p < 0.001, suggesting that the use of robustfit and exclusion of oscillatory peaks improves 1/f fitting (Kosciessa et al., 2020). There were no significant differences in RMSE values between fBOSC and eBOSC, p > 0.05.

**FIGURE 2.**
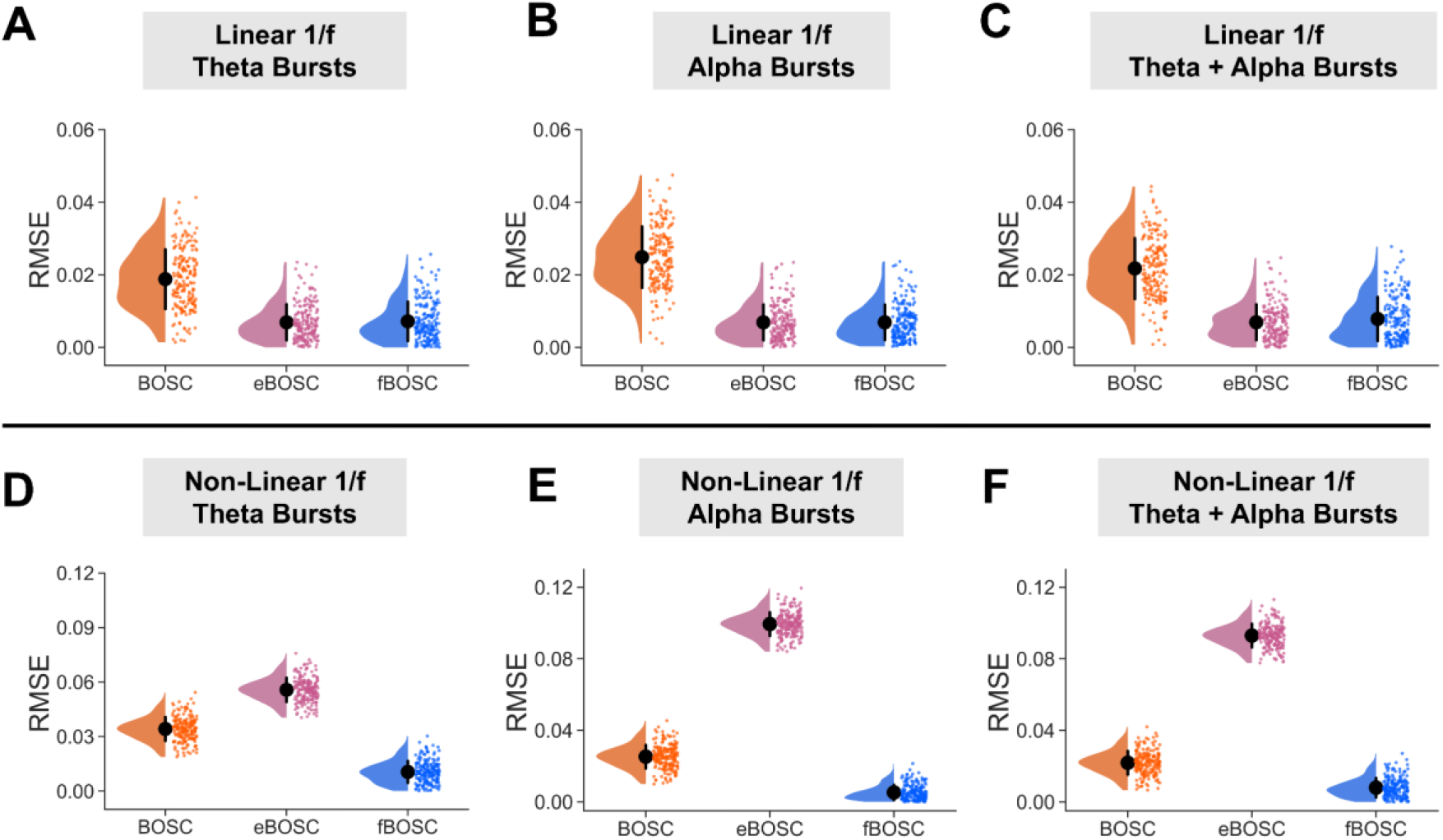
Simulations were performed using data with either a linear (A, B, C) or non-linear (D, E, F) background 1/f power spectrum. These data were then combined with simulated bursts in the theta-band alone (4 Hz), alpha-band alone (10 Hz), or both the theta- and alpha-bands (4 Hz and 10 Hz). The SNR of the bursts was varied between 5-12. For each set of simulations, the root mean squared error (RMSE) between the estimated and actual 1/f fit was plotted for BOSC, eBOSC and fBOSC. Individual data points correspond to RMSE values from each simulated trial. Error bars correspond to standard error.

Simulation results for data with a non-linear 1/f power spectrum are shown in Figure 2D-F. Across the different burst conditions (theta alone, alpha alone, theta and alpha) there was a main effect of method, F(2,597) > 2485, p < 0.001. Follow-up tests showed fBOSC had significantly lower RMSE values compared with the other two methods, all p < 0.001. RMSE values were especially large for eBOSC when theta frequencies were excluded from the 1/f fit (bottom panel, left and right). By excluding frequencies, the eBOSC 1/f fitting procedure assumes that the angle of the linear slope between 7-40 Hz extends below 7 Hz, when in fact there is a non-linear knee (also see Figure 1 middle panel). Consequently, eBOSC had higher RMSE values than fBOSC and BOSC for the simulated data with embedded theta bursts or a combination of theta and alpha bursts, p < 0.001. Similar results were obtained for simulations with very high SNR bursts (SNR varying between 24-48, see Supporting Figure S1).

When using fBOSC to model the simulated non-linear 1/f power spectrum, we chose to include an extra parameter, *k* (see Equation 1), to model the non-linear knee. This means that the model describing the 1/f fit for fBOSC was more complex than the models for BOSC and eBOSC. The lower RMSE values for fBOSC reported in Figure 2 may therefore have resulted from over-fitting rather than genuinely better modelling of the power spectrum. To investigate this further, we repeated the simulation analyses for the non-linear 1/f data and performed 10-fold cross validation (see Materials and Methods). For every fold and across all burst conditions (theta alone, alpha alone, alpha and theta), RMSE values for the test data were lowest for fBOSC, followed by BOSC and then eBOSC (Figure 3A-C). This suggests that using fBOSC with the knee option of FOOOF did not result in over-fitting and was the best of the three methods for modelling non-linear 1/f power spectra. We also performed another analysis to check for over-fitting – the simulation analyses were repeated using the AIC metric for model selection (see Materials and Methods). AIC values were lower for fBOSC than for BOSC and eBOSC (see Supporting Figure S2), again suggesting that despite being more complex, fBOSC performed background 1/f fitting better than BOSC and eBOSC, when power spectra were non-linear.

**FIGURE 3.**
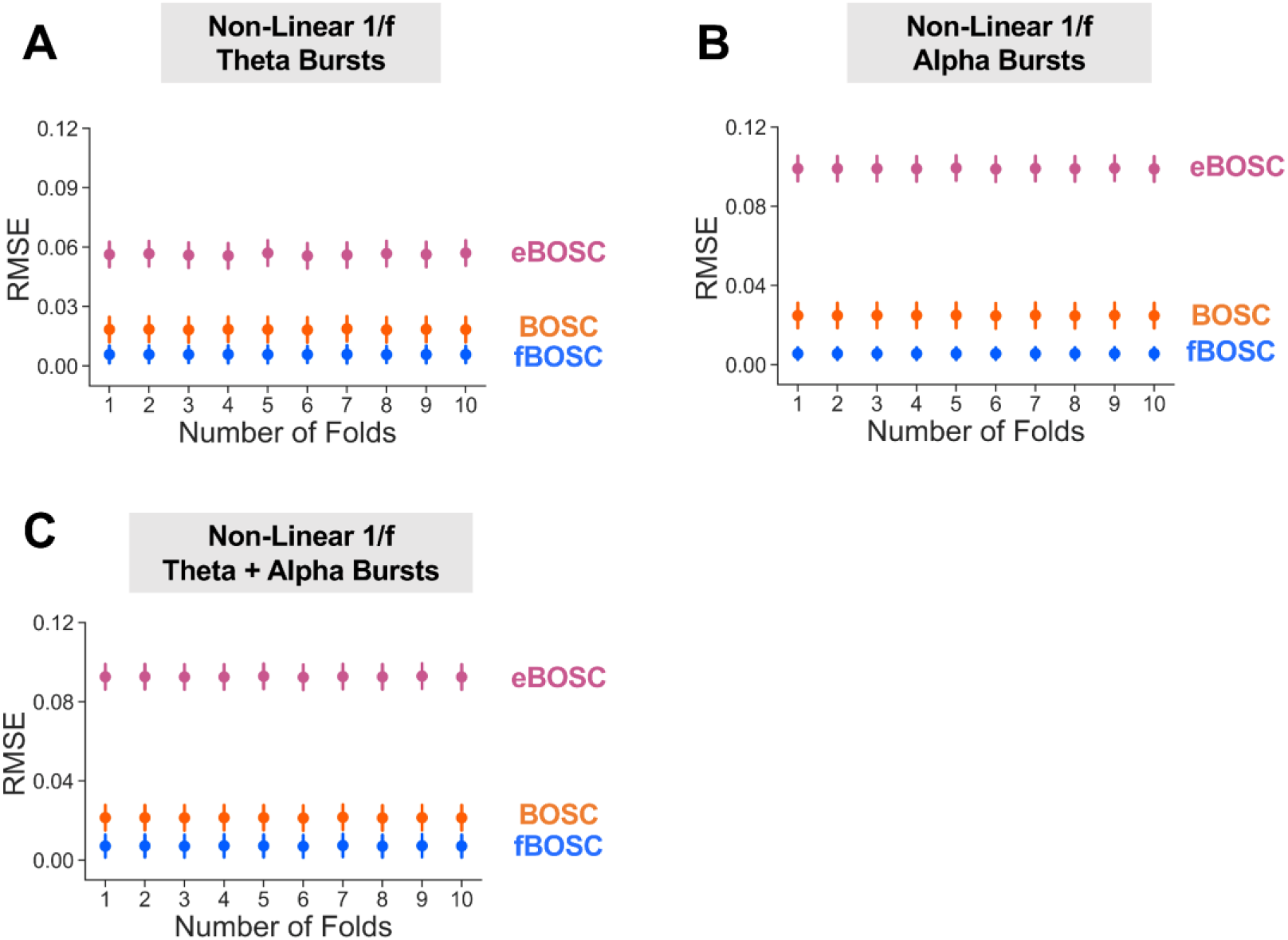
Ten-fold cross validation was performed using the simulated non-linear 1/f data with embedded oscillations in the theta-(A), alpha-(B) or theta- and alpha-bands (C). The RMSE of the test data is plotted for each method (BOSC, eBOSC and fBOSC) and for each fold. Error bars indicate standard error for each method and fold.

To assess how the three methods dealt with the presence of oscillatory peaks in the power spectrum, we repeated the simulation analyses using data with a 1/f non-linear background spectrum and oscillatory bursts in the theta- and alpha-bands. The SNR of the embedded bursts was increased from 0-24 in steps of 2, and no random variation in burst SNR was present as for the previous simulations. This had the effect of introducing increasingly large oscillatory peaks into the power spectrum. The RMSE between the simulated non-linear 1/f power spectrum and the fit from BOSC, eBOSC and fBOSC is plotted in Figure 4. eBOSC had the highest overall RMSE values, due to the exclusion of frequencies below 7 Hz when modelling the power spectrum. RMSE values also slightly increased as a function of SNR. BOSC had lower RMSE values than eBOSC, however these increased sharply as a function of SNR. This replicates a well-known issue with the BOSC background 1/f fitting approach (Kosciessa et al., 2020; Whitten et al., 2011). The RMSE error over trials was lowest for fBOSC, with a negligible increase as a function of oscillatory burst SNR. Overall, these simulations demonstrate that the 1/f fit from fBOSC is more accurate than the other two methods, including when the spectrum is non-linear and/or contains large oscillatory peaks.

**FIGURE 4.**
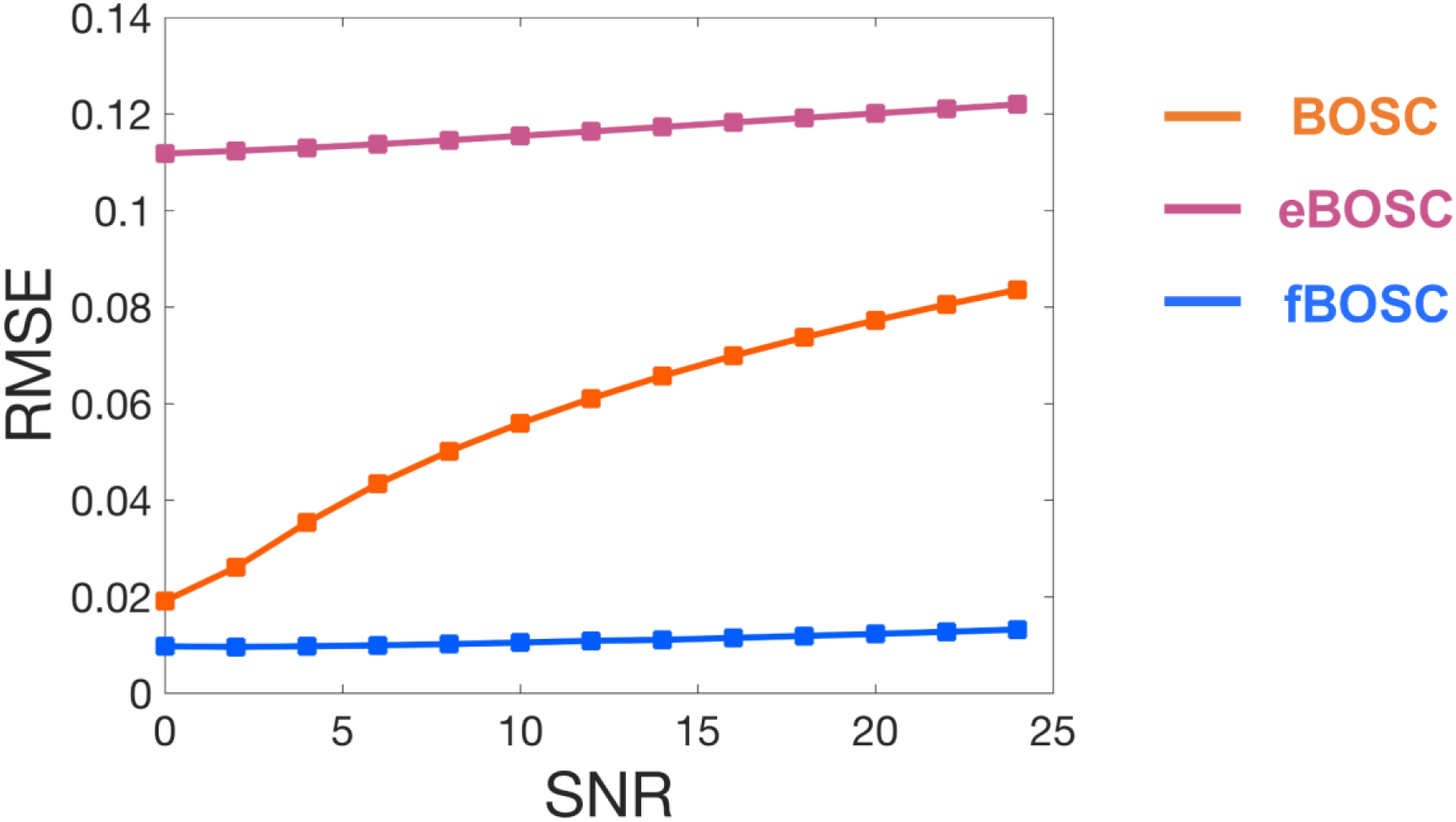
Simulations using data with a non-linear 1/f power spectrum and embedded theta and alpha bursts. The signal-to-noise ratio (SNR) of the bursts was increased from 0-24, in steps of 2. The root mean squared error (RMSE) between the estimated and actual 1/f fit was plotted for BOSC, eBOSC and fBOSC for each of the SNRs.

### fBOSC standardises burst detection across frequencies

Inaccuracies during the 1/f fit lead to inappropriate power thresholds for burst detection (see Figure 1, green line). In the case of non-linear power spectra, BOSC and eBOSC fail to model the knee, resulting in differing sensitives to oscillatory burst detection based on whether the frequency of the burst is above or below the knee, assuming a fixed power threshold is used. As an illustration of this, Supplementary Figure S3 shows an example theta burst detected using fBOSC but not eBOSC. To investigate this further, we performed a series of simulations with a non-linear 1/f background power spectrum, with embedded oscillatory bursts in theta-band (4 Hz) or alpha-band (10 Hz). The SNR of the bursts were scaled according to background 1/f activity at 4 Hz or 10 Hz. The hit rate and false alarm rates for theta/alpha burst detection were then compared between BOSC, eBOSC and fBOSC. As shown in Figure 5A, the hit rate for burst detection in the alpha-band was higher than the theta-band when using BOSC and eBOSC. However, for fBOSC, the hit rate was constant across theta and alpha (Figure 5A). Formal comparison of the difference between hit ratealpha and hit ratetheta showed that there was an effect of method F(2,597) = 129.4, p < 0.001, with fBOSC having significantly lower values than BOSC and eBOSC, p < 0.001. Unlike BOSC and eBOSC, the hit rate difference for fBOSC was close to 0 (Figure 5B), indicating no disparity between frequency bands.

**FIGURE 5.**
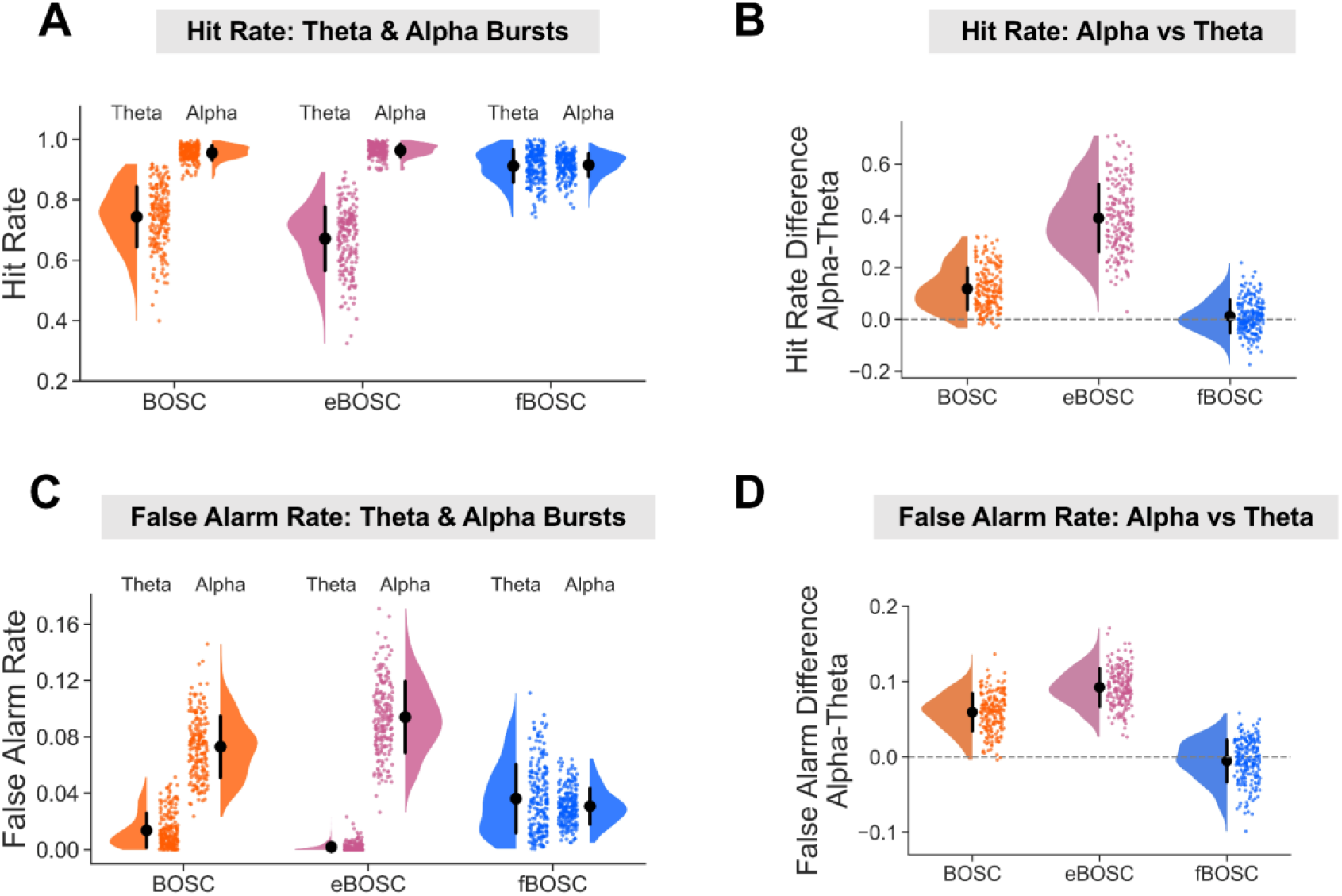
Simulations were performed using data with a non-linear 1/f power spectrum and embedded theta or alpha bursts (SNR varied between 5-12). BOSC, eBOSC and fBOSC were used to detect these oscillatory bursts. The (A) hit rate and (C) false alarm rate of theta and alpha burst detection are plotted for all three methods. The difference in (B) hit rate and (D) false alarm rate between theta and alpha burst detection is also plotted, with an additional dotted line at 0. Across all plots, individual data points correspond to each simulated trial. Error bars correspond to standard error.

False alarm rates were higher for alpha bursts than for theta bursts when using BOSC and eBOSC, but not for fBOSC (Figure 5C). Again, we subtracted false alarm ratealpha from false alarm ratetheta and compared the methods (Figure 5D). There was an effect of method F(2,597) = 174.0, p < 0.001, with fBOSC having lower values than the BOSC and eBOSC, p < 0.001. The difference in false alarm rate between alpha and theta bursts for fBOSC was close to 0 (Figure 4D). Similar false alarm results were obtained for simulations with very high amplitude bursts (SNR = 24-48, see Supporting Figure S4). Overall, our results show that when analysing data with a non-linear 1/f spectrum, fBOSC standardises the sensitivity for detecting oscillatory bursts across frequencies, in terms of hit rate and false alarm rate.

### Modelling 1/f activity: fixed versus knee parameter

The simulation analyses showed that fBOSC outperformed BOSC and eBOSC particularly in situations where the 1/f power-spectrum was non-linear. In human electrophysiological data, this non-linearity typically presents itself as a knee in the power spectrum from ∼0.5-10 Hz (Gao et al., 2017; He, 2014; He et al., 2010). To investigate whether modelling the knee reduces modelling errors in real data, we analysed MEG and iEEG resting state electrophysiological datasets (Larson-Prior et al., 2013; Miller et al., 2012; see Materials and Methods). The power spectrum from each ROI/electrode was computed using Welch’s method. Example power spectra are plotted from one MEG participant (Figure 6A) and one iEEG patient (Figure 6B). Note the non-linear nature of the spectrum when plotted in log-log space, with a prominent knee below ∼8 Hz. Across all MEG participants and iEEG patients the spectra were parametrised using the “fixed” option of the FOOOF algorithm (Donoghue et al., 2020b) which assumes a linear 1/f, or the knee option which assumes a non-linear 1/f. We specified a frequency range of 1-40 Hz, overlapping with the frequency of delta, theta, alpha and beta oscillations. To avoid over-fitting, we used 10-fold cross validation – 90% of the data was used as training data and parameterised using FOOOF (fixed and knee options). The resulting parameters were used to predict the power spectrum of the remaining held-out testing data, using RMSE to quantify performance. This was repeated for each fold of the data, and RMSE was averaged over each iteration. RMSE values were then compared between the fixed and knee options. Specifically, we subtracted RMSEfixed -RMSEknee. As shown in Figure 6, for both MEG (Figure 6C) and iEEG (Figure 6D) the difference in RMSE was significantly greater than 0, p < 0.001, suggesting increased accuracy when using the knee compared with the fixed option. This demonstrates how non-linear 1/f power spectra are common in human electrophysiological data when analysing frequencies between 1-40 Hz. We expect our fBOSC method to be particularly useful in these situations, and especially when analysing oscillatory bursts overlapping with the knee frequency (∼0.5-10 Hz).

**FIGURE 6.**
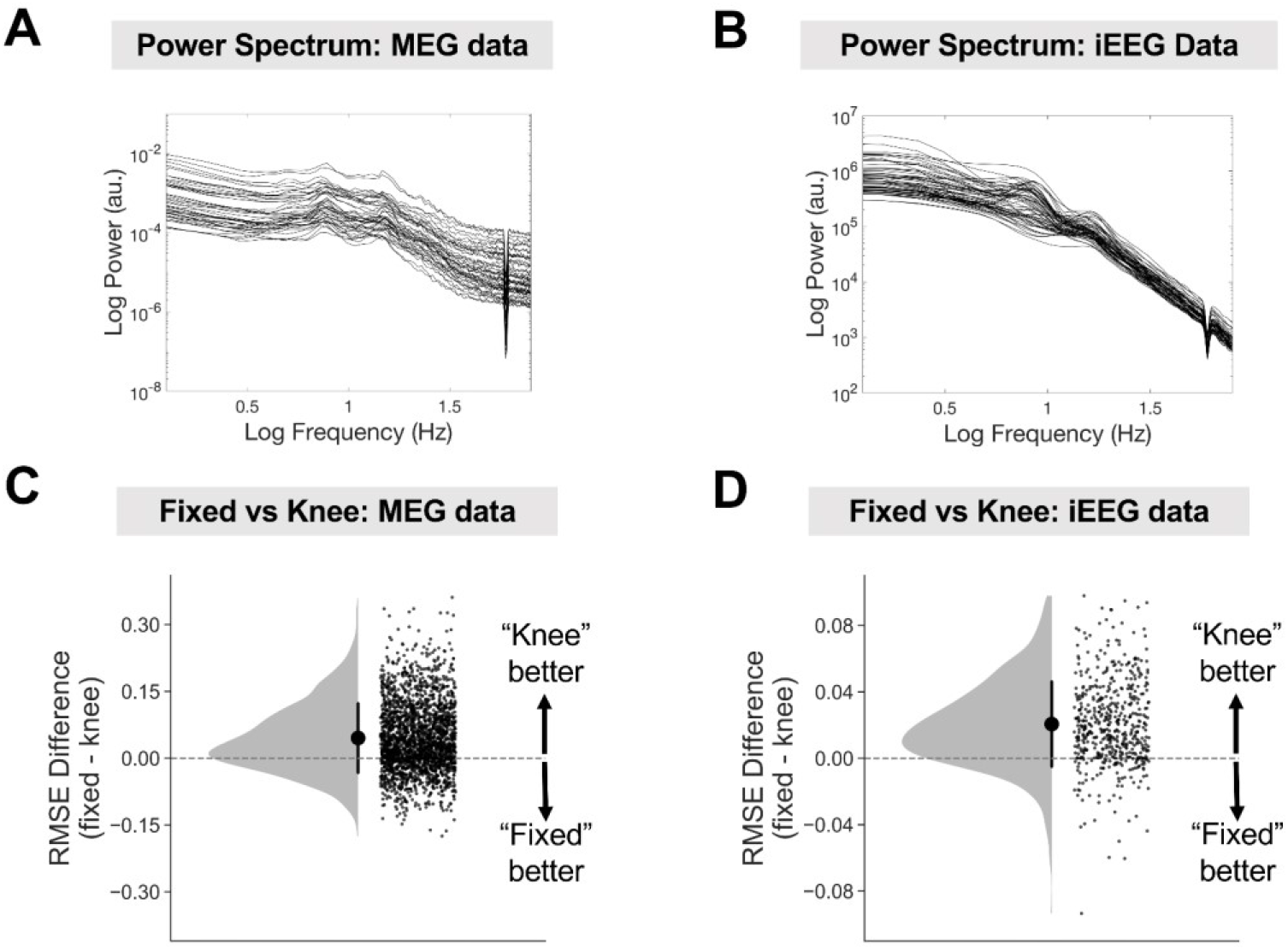
Neural power spectra are often non-linear. Power spectra are plotted on a log-log scale from one example MEG dataset (A) and one iEEG dataset (B). Individual lines correspond to ROIs (MEG) or electrodes (iEEG). Note the prominent knee in the power spectrum below ∼8 Hz. The difference in RMSE when modelling the 1/f power spectrum as linear (fixed) or non-linear (can contain a knee parameter) using FOOOF (Donoghue et al., 2020b) was plotted for MEG (C) and iEEG (D) datasets. Individual data points correspond to ROIs across participants (MEG) or electrodes across patients (iEEG). Error bars correspond to standard error.

### fBOSC and theta-band burst detection

Next, we used BOSC, eBOSC and fBOSC to detect oscillatory bursts in the same MEG and iEEG datasets with the aim of quantifying differences in burst detection between the methods. We focussed on theta-band (3-7 Hz) bursts where errors in the 1/f fit will be highest for non-linear power spectra when using BOSC or eBOSC (Figure 1). Quantifying the bursting properties of theta rhythms is of particular interest for the study of working memory (Lisman, 2010), autobiographical memory retrieval (Barry et al., 2019) and spatial navigation in humans (Stangl et al., 2021).

For the MEG dataset (Larson-Prior et al., 2013), across 50 participants we quantified theta burst “abundance”, defined as the duration of the theta rhythmic episode relative to the length of the analysed data segment. Abundance values are scaled between 0 and 1, where 0 indicates no burst present and 1 indicates a burst continuously present. This is crucial metric in burst analysis as it helps separate rhythmic duration from power (Kosciessa et al., 2020). Focussing on the ROI with the highest theta abundance (right dorsolateral prefrontal cortex, Figure 7A), we statistically compared abundance values as quantified using BOSC, eBOSC and fBOSC (Figure 7B). There was a main effect of method, *F*(2,147) = 8.151, p < 0.001, with follow-up tests showing that fBOSC produced higher theta abundance values than the other two methods, p < 0.005. This result is unsurprising, given that the 1/f fit will be lower at theta frequencies for fBOSC than for BOSC and eBOSC, accurately reflecting the knee in the power spectrum (Donoghue et al., 2020b; He, 2014; He et al., 2010). We also quantified the mean SNR of theta bursts versus the background spectrum at 3-7 Hz. There was a main effect of method, *F*(2,147) = 46.77, p < 0.001, with follow-up tests showing a reduction in mean theta SNR for fBOSC versus the other two methods, p<.001 (Figure 7C).

**FIGURE 7.**
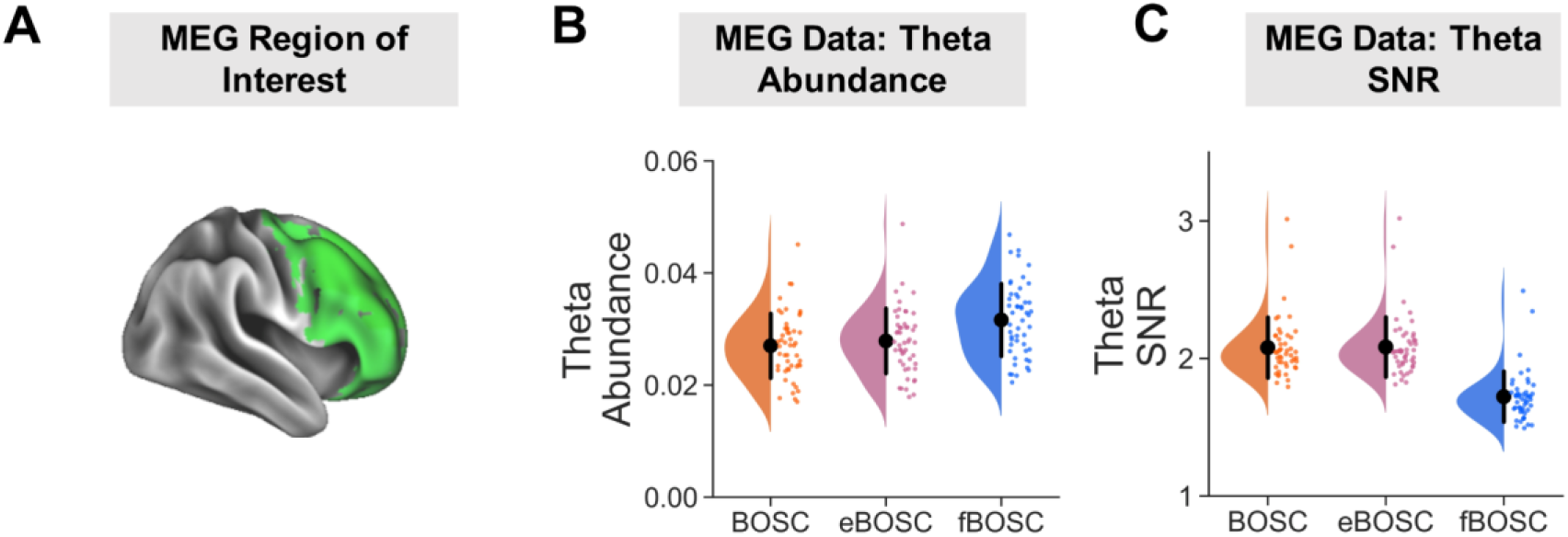
BOSC, eBOSC and fBOSC were used to detect oscillatory bursts in resting state MEG data. (A) We focussed on a region of interest in the right dorsolateral prefrontal cortex, where theta bursts were most prominent (vertices within the ROI are plotted in green and rendered on a cortical mesh using Connectome Workbench). (B) Theta (3-7 Hz) burst abundance was quantified as the duration of rhythmic episodes relative to the length of the recording and plotted separately for each method. (C) Theta burst SNR was quantified as the power of each burst relative to the background 1/f fit at 3-7 Hz. Individual data points correspond to each MEG participant. Error bars correspond to standard error.

For the iEEG dataset (Miller et al., 2012), we repeated the theta abundance analysis using data from 533 electrodes across 10 participants (see Materials and Methods). Theta abundance values were statistically compared using BOSC, eBOSC and fBOSC (Figure 8A). Again, there was a main effect of method, F(2,1596) = 57.43, p < 0.001, with follow up tests showing that fBOSC produced higher theta abundance values than BOSC and eBOSC, p < 0.001. The SNR of each theta burst was also quantified using all three methods (Figure 8B). There was a main effect of method, F(2,1596) = 7.53, p < 0.001, with follow up tests showing that fBOSC produced higher SNR values than eBOSC, p < 0.001, and there was a trend for higher fBOSC SNR values compared with BOSC, p = .053. Overall, our analysis of theta bursts in human MEG and iEEG electrophysiological datasets showed that using fBOSC produced statistically different results compared to BOSC or eBOSC. Specifically, at a set fixed threshold, BOSC and eBOSC produced lower theta abundance and either lower (MEG) or higher (iEEG) SNR values when a knee was present in the power spectrum.

**FIGURE 8.**
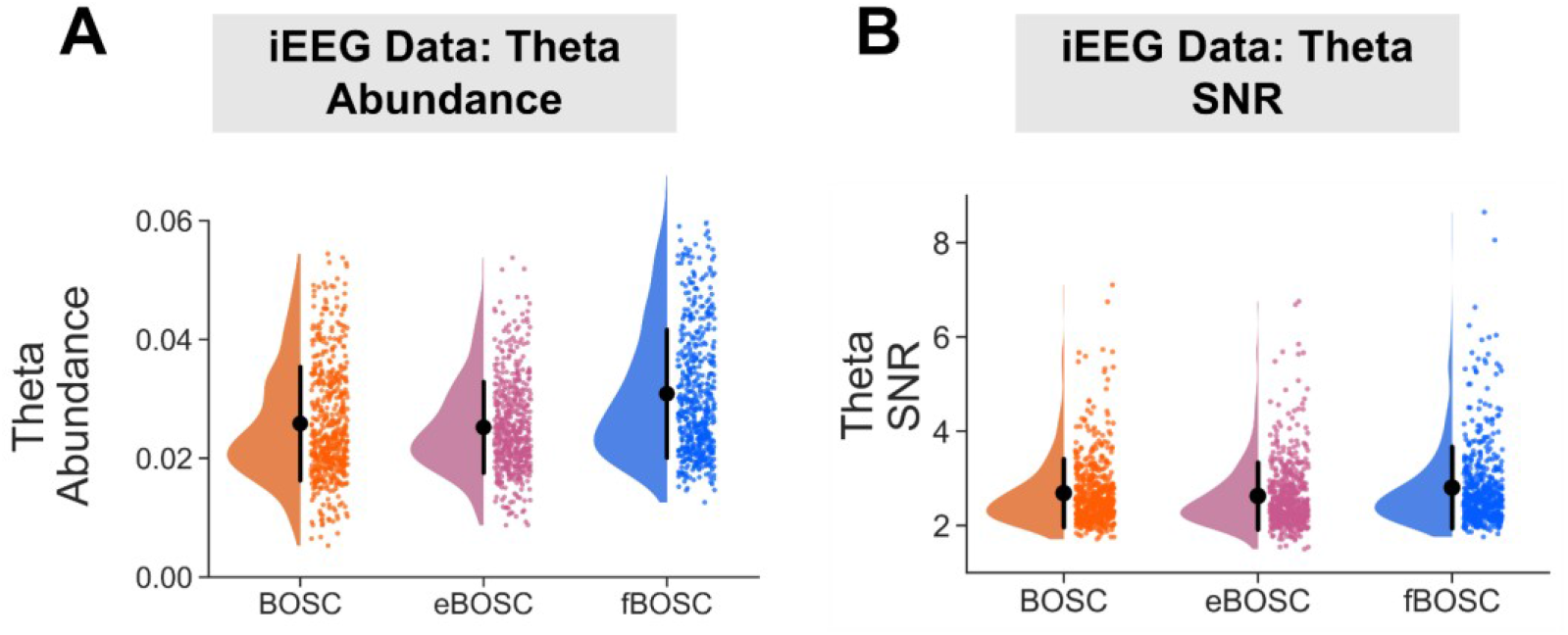
Using resting state iEEG data from 10 patients, theta (3-7 Hz) burst abundance (A) was quantified as the duration of rhythmic episodes relative to the length of the recording. (B) Theta burst SNR was quantified as power of each burst relative to background 1/f fit at 3-7 Hz. Abundance and SNR values are plotted separately for each method. Individual data points correspond to each of the 533 electrodes across the 10 patients. Error bars correspond to standard error.

## DISCUSSION

In this study we presented an improved method for oscillatory burst detection based on the BOSC framework (Caplan et al., 2001; Hughes et al., 2012). To separate background 1/f neural activity from rhythmical bursts, the BOSC framework models the average neural power spectrum across trials to define a power threshold per frequency of interest. Rather than using existing linear regression approaches, here we utilised a recently developed spectral parametrisation algorithm (Donoghue et al., 2020b), which accurately models neural power spectra across a wide variety of conditions.

A series of simulation analyses were performed to compare our modified method, fBOSC, with existing approaches: (i) the original BOSC implementation which uses a partial least squares regression for 1/f fitting, and (ii) the extended BOSC implementation (Kosciessa et al., 2020) which uses MATLAB’s robustfit function for 1/f fitting combined with manual removal of oscillatory peaks (Figure 1). Simulation analyses for data with a linear power spectrum showed that fBOSC more accurately modelled the 1/f slope than the original BOSC implementation, and performed similarly to eBOSC. We also replicated an established issue with the original BOSC implementation (Whitten et al., 2011), whereby the fit is biased by peaks in the power spectrum. Simulations were also performed using data with a non-linear 1/f slope, containing a prominent knee below ∼5 Hz (Gao et al., 2020; He, 2014). Both BOSC and eBOSC failed to accurately model the power spectrum in this context resulting in high RMSE values. The errors were particularly high for eBOSC after removal of peaks below the knee frequency. These results are unsurprising, given that both methods assume a linear 1/f slope. By contrast FOOOF can model neural power with an additional knee parameter and, consequently, across all simulations, fBOSC produced the lowest modelling errors (Figure 2). These results were not caused by over-fitting from the addition of the extra knee parameter, k (Figure 3 and Supporting Figure S2). Furthermore, fBOSC was shown to be relatively unaffected by the presence of increasingly large oscillatory peaks in the power spectrum (Figure 4). This is presumably because FOOOF models peaks as gaussians, iteratively removes them from the power spectrum, and then re-models the flattened 1/f slope (Donoghue et al., 2020b). This leads to more accurate 1/f fits compared with the exclusion of peaks as used in eBOSC (Kosciessa et al., 2020). We also note that the modelling and removal of peaks is automated through the use of FOOOF, whereas the exclusion of peaks requires user specification with eBOSC.

Within the BOSC framework, the 1/f fit is used to directly determine the power threshold. Consequently, any inaccuracies introduced during the 1/f modelling process will have knock-on effects for burst detection. Where oscillatory peaks bias the 1/f fit (Figure 4), the power threshold will be artificially increased across some or all frequency bands of interest. The failure to model the knee in the power spectrum has even greater consequences, as shown in Figure 5. Using simulated data with a non-linear 1/f power spectra, BOSC and eBOSC had dramatically different hit rates and false alarm rates between theta-band and alpha-band bursts. This is because theta bursts occur below the knee frequency, whereas alpha bursts occur above the knee frequency. Of course, this assumes a fixed threshold was used across frequencies (e.g. the 99^th^ percentile of the theoretical probability distribution). In contrast, fBOSC displayed identical hit rate and false alarm rates for theta and alpha bursts embedded within simulated data with a non-linear background 1/f spectrum. Our fBOSC method, therefore, ensures equivalent sensitivity for burst detection across frequency bands.

The empirical application of fBOSC was demonstrated using MEG and iEEG resting state datasets, which showed evidence of non-linear power spectra (Figure 6). We focussed on quantifying theta-band (3-7 Hz) bursts below the ∼5 Hz knee. There were quantifiable differences in the abundance and SNR of theta bursts when using fBOSC compared to BOSC or eBOSC. Abundance values were approximately 1% higher for the iEEG data when using fBOSC (Figure 8). This is presumably due to the failure of BOSC and eBOSC to accurately model the bend in the power spectrum overlapping with theta frequencies. Interestingly, theta SNR burst increased when using fBOSC for the iEEG dataset, but decreased for the MEG dataset, highlighting the dissociation between burst abundance and amplitude between datasets (Kosciessa et al., 2020). On a practical level, fBOSC is expected to give the greatest benefits over and above existing BOSC methods for the detection of bursts below or overlapping with the knee frequency, i.e. delta (1-2 Hz), theta (3-7 Hz) and potentially alpha (8-13 Hz) bursts. This could be particularly useful in the field of memory research to disentangle the roles of theta oscillations from dynamic tilts in the background 1/f spectrum (Herweg et al., 2020). In terms of higher-frequency bursts (e.g. beta, gamma), the advantages of using fBOSC compared with the existing methods are likely to be subtler. In neural data, frequencies from 10-100 Hz are accurately modelled with linear approaches (He, 2014; Kosciessa et al., 2020; Miller et al., 2012), although our simulation analyses (Figure 2, top panel) did reveal that fBOSC significantly improved 1/f fitting for linear power spectra compared to BOSC.

More generally, our findings highlight the importance of robustly separating background 1/f activity from rhythmical activity. Neural oscillations are generated by groups of neuronal ensembles firing in a regular, synchronised manner (Buzsaki, 2006; Buzsáki & Draguhn, 2004). These often occur within single trials as transient bursts (Bonaiuto et al., 2021; Jones, 2016; Stokes & Spaak, 2016). On the other hand, background 1/f neural activity is highly correlated with asynchronous population neuronal firing rates in macaques and humans (Manning et al., 2009). Interestingly, the knee frequency at 0.5-10 Hz is related to a decay constant (Gao et al., 2020), potentially from membrane leak (Miller et al., 2009). Rhythmical oscillations and arrhythmical 1/f activity are concurrently measured with iEEG/EEG/MEG, but are clearly dissociable in terms of their neural origins (He, 2014; Miller et al., 2012). Both types of activity are physiologically important for cognition, but conflating them has consequences for the interpretation of neuroscientific findings. There is emerging evidence that previously reported oscillation-related effects might actually be driven by a spectral tilt of the 1/f power spectrum (Donoghue et al., 2020b; Herweg et al., 2020; Lendner et al., 2020; Ouyang et al., 2020). For example, He et al. (2019) recently reported that the classic developmental redistribution of oscillatory power from lower to higher frequencies during childhood can be partly explained by a flatter 1/f slope in children compared to adults. Calculating the power ratio between frequency bands also conflates rhythmical and 1/f arrhythmical components of neural signals (Donoghue et al., 2020a).

One of the primary goals of any burst detection algorithm should be the robust isolation of rhythmical signals from background 1/f activity and other experimental noise (Donoghue et al., 2021; van Ede et al., 2018). fBOSC achieves this with greater accuracy and flexibility than existing BOSC methods. In addition, fBOSC returns parameters describing the shape of background 1/f activity from the FOOOF parametrisation (offset, exponent and knee), which can be examined separately from any oscillatory bursting properties. These parameters could, for example, be used to estimate population neuronal firing rates between participants (Voytek et al., 2015), between different experimental conditions (Gao et al., 2020) or between different stages of sleep (Lendner et al., 2020).

When analysing neural power spectra, there are several methodological considerations to be considered. We refer the interested reader to Donoghue et al. (2020b) and Gerster et al. (2021) for thorough guidelines. One commonly encountered issue is the modelling of spectra with oscillatory peaks crossing the edge of the frequency range. This creates large fitting errors as FOOOF models complete gaussian peaks (Gerster et al., 2021). If the situation is unavoidable, users should specify a smaller fitting range at higher frequencies (via the `cfg.fBOSC.fooof.settings` option). One particularly important user-defined parameter when using FOOOF is whether to model the power spectrum with (knee) or without (fixed) a non-linear bend. Generally, this should be present in human neurophysiological data around 0.5-10 Hz (Chaudhuri et al., 2017; Gao et al., 2020). We opted to use a cross validation procedure to quantify modelling errors when using FOOOF with the knee or fixed options for MEG and iEEG datasets between 1-40 Hz, concluding that it was better to use the knee option. A similar procedure could be used for model selection in other datasets to determine whether modelling the knee is warranted.

More generally, fBOSC users should routinely visualise the 1/f fit using `fBOSC_fooof_plot.m` to identify inaccurate 1/f fitting results. When performing burst analysis, we would also recommend estimating the background 1/f spectrum separately between experimental conditions or participant groups of interest. This ensures that any reported burst-related differences are not simply a reflection of a spectral tilt of the 1/f slope between conditions or groups (He et al., 2019; Weber et al., 2020; Wilson et al., 2022). The parametrised 1/f slope properties returned by fBOSC could also be compared between conditions. It should be noted that FOOOF (Donoghue et al., 2020b) is not the only spectral parametrisation tool available. One popular alternative is Irregular-Resampling AutoSpectral Analysis (IRASA) which estimates fractal activity through a resampling procedure (Wen & Liu, 2016). However, IRASA operates in the time-domain rather than on power spectra, and can distort 1/f fits when data contain a knee (Donoghue et al., 2020b). FOOOF also has reduced computational costs compared with IRASA.

The BOSC framework, which detects bursts via amplitude and duration thresholds, is an intuitive, computationally inexpensive and flexible tool for oscillatory burst analysis (Caplan et al., 2001; Hughes et al., 2012). It exists alongside a plethora of other burst detection techniques. Our method, fBOSC, is conceptually similar to the recently developed Periodic/Aperiodic Parameterization of Transient Oscillations (PAPTO) approach (Brady & Bardouille, 2022), in that background 1/f activity is parametrised using FOOOF (Donoghue et al., 2020b). However, PAPTO has a different amplitude threshold procedure (Shin et al., 2017) and post-processing options. Other methods include the use of Hidden Markov Models (HMMs) to characterise transient changes in spectral power across multiple frequency bands (Quinn et al., 2019). However, it is unclear whether HMM-based methods are able to properly separate dynamically changing 1/f activity from oscillatory bursting across frequencies. It is also important to note that where neural oscillations are non-sinusoidal or possess some other non-linear property, the Fourier-based decomposition of signals used within the BOSC framework will be unsuitable and may lead to spurious results (Donoghue et al., 2021b). Non-sinusoidal oscillatory bursts would be better quantified through empirical mode decomposition (Huang et al., 1998; Quinn et al., 2021) or time domain approaches based on waveform shape (Cole & Voytek, 2019). A principled comparison between different burst detection methods is beyond the scope of this article, but would be of benefit to the field.

One way in which fBOSC could be improved upon in the future is in terms of the dynamic estimation of background activity. In the current implementation, time-frequency spectra are averaged with the assumption that 1/f activity is constant across trials or conditions. However, it is well established that background 1/f activity can be dynamic, changing with arousal level (Lendner et al., 2020) and during cognitive tasks (Gao et al., 2020). Time varying spectral parametrisation approaches (Wilson et al., 2022) are in development, and could be used to dynamically update the power threshold used for burst detection. This would be particularly useful for situations where dynamic 1/f changes co-occur with oscillatory bursts, as well as for longer EEG/MEG sleep recordings where the tilt of the 1/f slope changes dramatically between sleep stages (Lendner et al., 2020). Of course, this would come with the added computational effort of parametrising multiple neural spectra.

## Conclusions

We have presented a tool for oscillatory burst detection which combines spectral parametrisation (FOOOF, Donohough et al., 2020b), with the BOSC framework, termed fBOSC. This modification addresses two issues with existing methods when modelling the 1/f background spectrum of neural data (Caplan et al., 2001; Kosciessa et al., 2020; Whitten et al., 2011). First, it is robust to oscillatory peaks in the power spectrum. Second, it can accurately model non-linear power spectra containing a knee. By robustly separating background 1/f activity from neural oscillations, fBOSC ensures that the power threshold used for burst detection is consistent across frequencies. Our tool is openly available for use at https://github.com/neurofractal/fBOSC.

## Abbreviations

AIC: Akaike information criterion
BOSC: better oscillation
EEG: electroencephalography
FOOOF: fitting oscillations and one over f
HMM: hidden Markov model
iEEG: intracranial electroencephalography
MEG: magnetoencephalography
PAPTO: Periodic/Aperiodic parameterization of transient oscillations
RMSE: root mean squared error
SNR: signal-to-noise ratio

## ACKNOWLEDGEMENTS

This research was supported by a Wellcome Principal Research Fellowship to E.A.M. (210567/Z/18/Z). Thanks to Gareth Barnes and the UCL OP-MEG team for helpful discussions throughout the development of fBOSC. We also thank Jeremy Caplan, Adam Hughes, Tara Whitten (BOSC developers) as well as Julian Kosciessa and colleagues (eBOSC developers), for making their code openly available.

This research was funded in whole, or in part, by Wellcome (Grant number: 210567/Z/18/Z). For the purpose of Open Access, the authors have applied a CC BY public copyright licence to any Author Accepted Manuscript version arising from this submission.

## AUTHOR CONTRIBUTIONS

**Robert A. Seymour:** Conceptualisation, Methodology, Software, Formal Analysis, Discussion, Writing – Original Draft; **Nicholas Alexander:** Discussion, Writing – Review and Editing; **Eleanor A. Maguire:** Supervision, Funding Acquisition, Discussion, Writing – Review and Editing.

## DATA SHARING AND DATA AVAILABILITY

Data and code for reproducing our simulation results in Figures 2-5 can be found at https://github.com/neurofractal/fBOSC_validation. The pre-processed resting state MEG data used in this study are openly available for download from the Human Connectome Project (https://db.humanconnectome.org/; Larson-Prior et al., 2013). The iEEG resting state data are also openly available for download (https://purl.stanford.edu/zk881ps0522; fixation_PAC.zip; Miller et al., 2012).

## SUPPORTING INFORMATION

Is available -Supporting Figures S1, S2, S3 and S4.

## Supporting Information

**SUPPORTING FIGURE S1.**
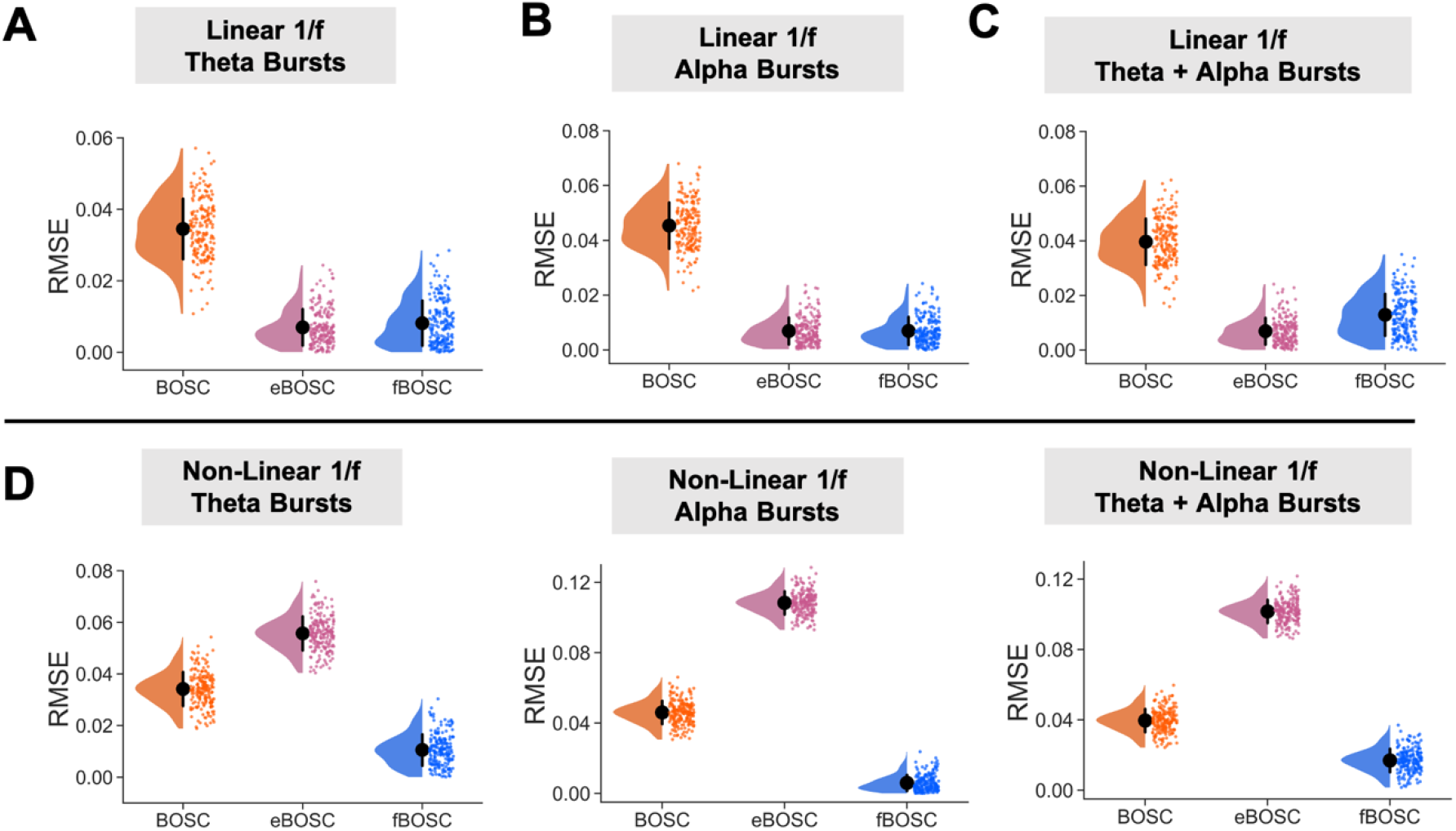
Simulations were performed using data with either a linear (A-C) or non-linear background 1/f power spectrum. These data were then combined with simulated bursts in the: theta-band (4 Hz) alone; alpha-band (10 Hz) alone; or both the theta and alpha-bands (4 Hz and 10 Hz). The SNR of the bursts was varied between 24-48. For each set of simulations, the root mean squared error (RMSE) between the estimated and actual 1/f fit was plotted for BOSC, eBOSC and fBOSC. Individual data points correspond to RMSE values from each simulated trial. Error bars correspond to standard error. The pattern of results is very similar to those shown in main Figure 2 (where the SNR of bursts varied between 5-12).

**SUPPORTING FIGURE S2.**
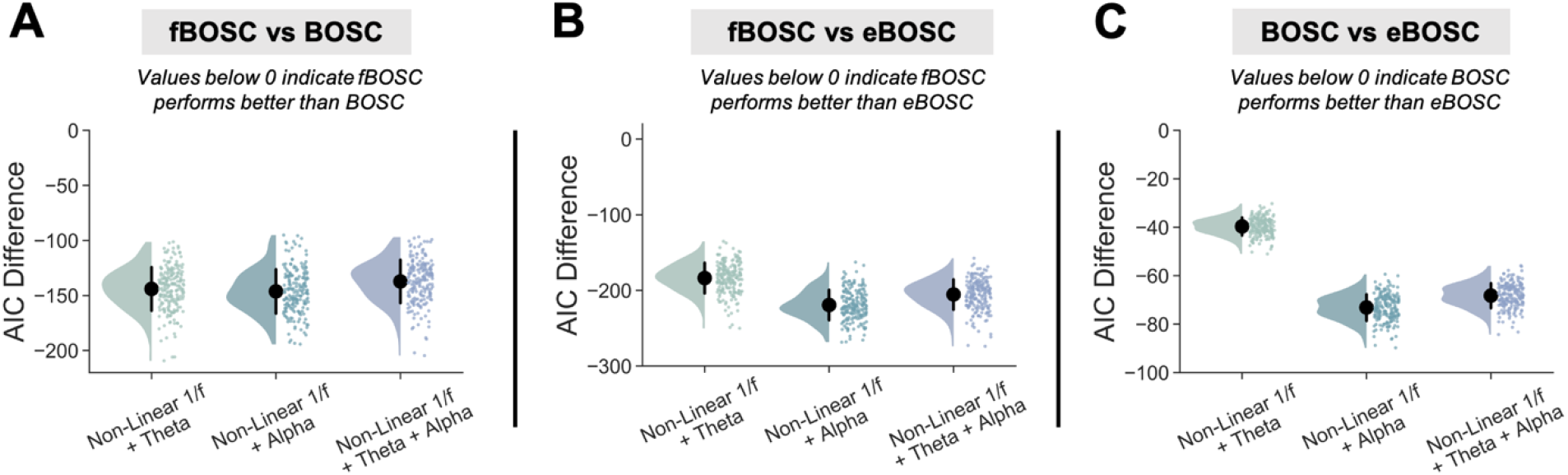
Simulations were performed using data with a non-linear 1/f power spectrum and embedded oscillations in the theta-, alpha-or theta- and alpha-bands. The error between the estimated and actual 1/f fit was quantified using the Akaike information criterion (AIC) for each method (BOSC, eBOSC and fBOSC). For model comparison, AIC values were compared between fBOSC and BOSC (A); fBOSC versus eBOSC (B); and between BOSC and eBOSC (C). Error bars indicate standard error for each method, and individual datapoints correspond to each simulated trial. Note that fBOSC has lower AIC values than the other two methods, indicating better accuracy at recovering the non-linear background 1/f activity.

**SUPPORTING FIGURE S3.**
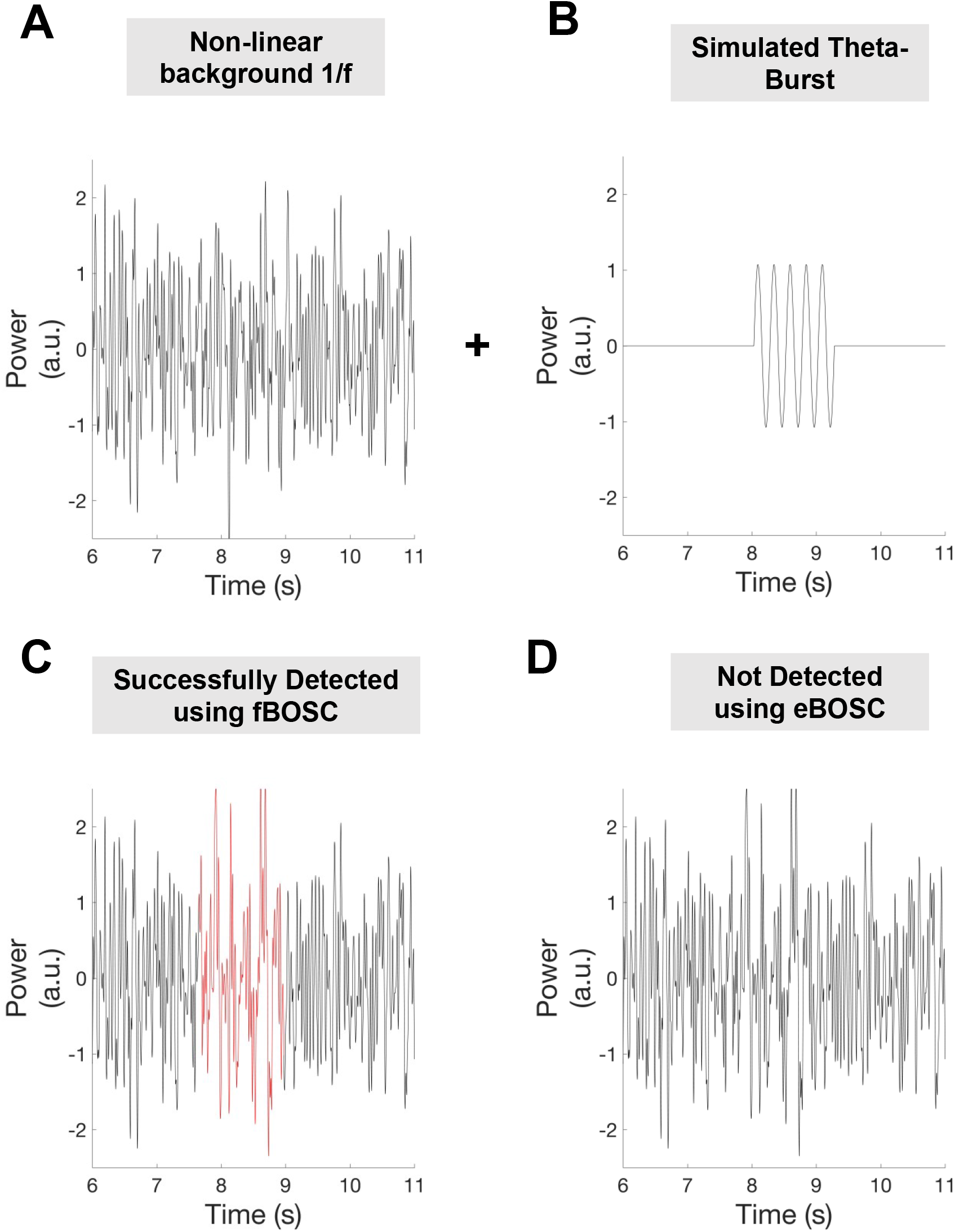
An example theta burst detected by fBOSC but not eBOSC. A non-linear 1/f spectrum was simulated (A) and combined with a simulated burst at 4 Hz (SNR = 9, compared with the background spectrum at 4 Hz) from ∼ 8 s to 9 s in this dataset (B). Burst detection was performed using fBOSC and eBOSC using the same parameters as specified in the main manuscript. Detected burst times are plotted in red. fBOSC detected this burst (C) but eBOSC did not (D).

**SUPPORTING FIGURE S4.**
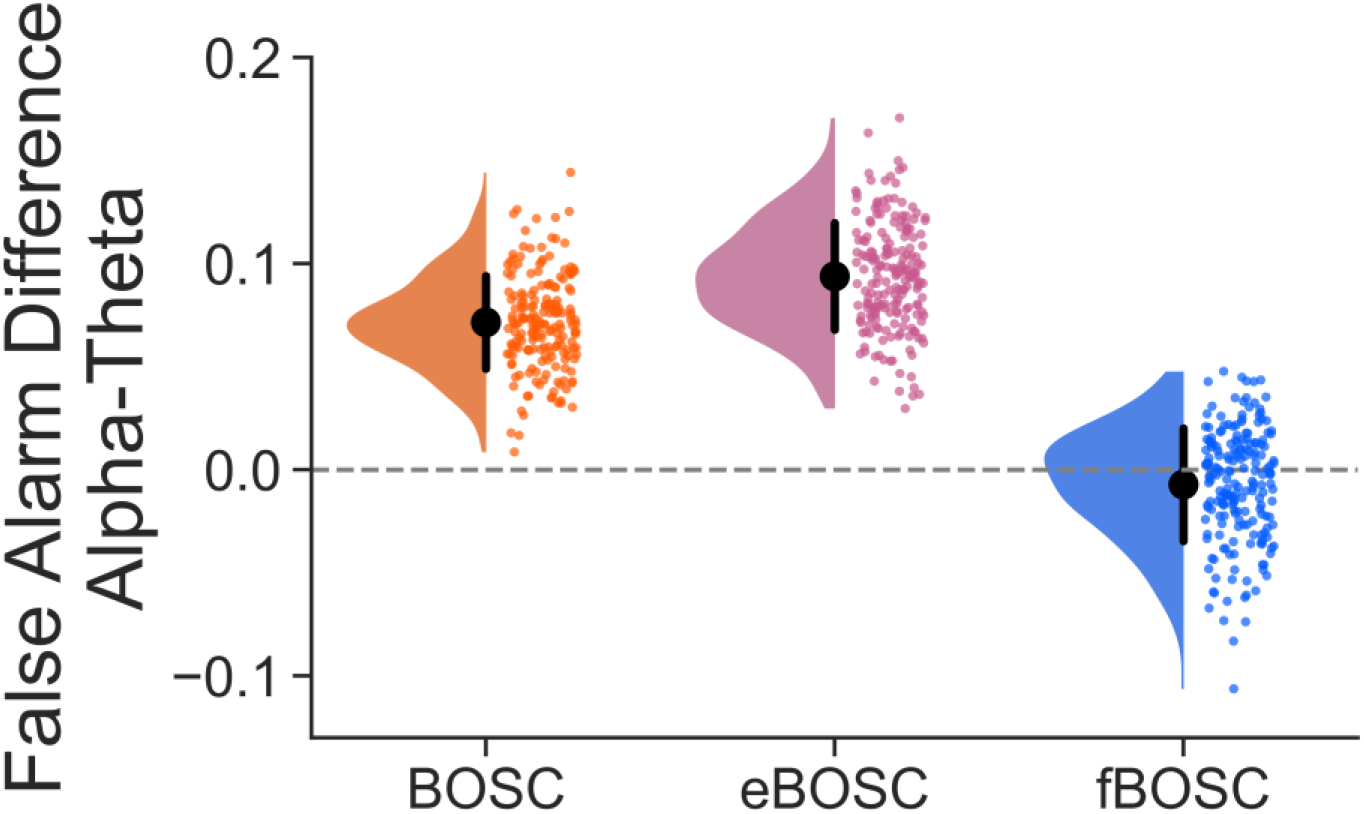
Simulations were performed using data with a non-linear 1/f power spectrum and embedded theta or alpha bursts (the SNR of bursts varied between 24-48). BOSC, eBOSC and fBOSC were used to detect these oscillatory bursts. The difference in false alarm rate between theta and alpha burst detection is plotted, with an additional dotted line at 0. Across all plots, individual data points correspond to each simulated trial. Error bars correspond to standard error. Note that fBOSC is much closer to 0 than the other two methods, indicating smaller differences in the false alarm rate between frequency bands. Hit rate is not plotted because the high SNR bursts caused ceiling effects, i.e. the hit rate was nearly always equal to 1 for each trial.

